# Yeast extract improves growth in rainbow trout (*Oncorhynchus mykiss*) fed a fishmeal-free diet and modulates the hepatic and distal intestine transcriptomic profile

**DOI:** 10.1101/2023.02.23.529675

**Authors:** Laura Frohn, Diogo Peixoto, Cervin Guyomar, Carla Teixeira, Frédéric Terrier, Pierre Aguirre, Sarah Maman Haddad, Julien Bobe, Benjamin Costas, Nadège Richard, Karine Pinel, Sandrine Skiba-Cassy

## Abstract

Replacing fishmeal with alternative protein sources and improving new ingredients diets with feed additives are major objectives in aquaculture. The aim of this study was to evaluate benefits for rainbow trout (*Oncorhynchus mykiss*) of supplementing a fishmeal-free diet, composed of processed animal proteins, with yeast extract. Juvenile rainbow trout (initial weight 37 ± 2 g) were fed either with a control diet (19% fishmeal) or with a diet based on terrestrial animal by-products (17%) supplemented or not with 3% of yeast extract. Effects of the diets were evaluated in a 4-week digestibility trial and a 12-week growth experiment. Fish health was investigated by measuring plasma immune markers and performing histological study of the gut. Underlying molecular responses were investigated using unbiased transcriptomic analysis of the liver and distal intestine. Results indicated that supplementing with 3% yeast extract did not influence nutrient digestibility substantially. Nevertheless, fish fed the supplemented fishmeal-free diet grew more than those fed the non-supplemented processed animal protein diet. Plasma and structural parameters indicated no exacerbated immune response or signs of intestinal inflammation in fish fed the fishmeal-free diets. However, plasma total immunoglobulin M levels and intestinal villi were significantly higher in fish fed the diet supplemented with yeast extract. The transcriptomic analysis revealed that the diets influenced immune, inflammatory, pathogen fighting and coagulation gene-related expressions. These results suggest that the dietary inclusion of yeast can enhance a fishmeal-free diet by improving rainbow trout performances and potentially their robustness.

## 1. Introduction

According to the most recent forecasts of the Food and Agriculture Organization, global protein consumption will increase by 14% by 2030. The production of aquatic protein will help meet the global demand for protein, since per capita fish consumption is expected to exceed more than 21 kg by then. Fishing and aquaculture activities meet financial and food requirements for populations around the world. Both sectors together employ nearly 8% of the world’s population and provide many people with a major source of dietary protein. In 2020, 49% of the 178 million tons of total fish production came from aquaculture and the sector is expected to contribute more than 50% of total production by the end of the decade. Thus, aquaculture is recognized as a major contributor to global food security ^(1,2)^.

Although most of the production is intended for direct human consumption, some is used for other purposes. Currently, 12% of the total annual fish tonnage is used to produce fish oil and fishmeal to feed carnivorous farmed fish such as Atlantic salmon (*Salmo salar*) and rainbow trout (*Oncorhynchus mykiss*) ^(1)^. These ingredients have been used to formulate aquafeeds since they fulfill nutritional needs for protein, amino acids and n-3 long-chain polyunsaturated fatty acids, and they are highly palatable and digestible ^(3)^. Because wild fish stocks used for this purpose have been overexploited for years, they are now protected by fishing quotas to conserve them and their use in carnivorous fish feeds tends to decrease ^(4)^.

Thus, alternative ingredients are increasingly of interest and many studies on the topic have emerged in the past few decades. For salmonids, changes in formulations have enabled replacing a large percentage of marine ingredients with plants raw materials ^(5)^. However, the presence of anti-nutritional factors and an imbalance in amino acids prohibit completely replacing fishmeal with plants since doing so decreases fish growth ^(6,7)^. Moreover, certain abrasive plant-based ingredients, especially soybean meal, trigger enteritis and consequently weaken the immune response in fish ^(8)^. Insect meal and microalgae are interesting alternatives due to their high nutritional qualities and the short life cycles of the organisms that provide the ingredients. However, using them as a complete substitute for fishmeal in aquafeeds is not recommended, since they are more expensive and raise concerns about environmental sustainability ^(9,10)^. Processed animal proteins (PAP), banned from Europe in 2001 due to spongiform encephalopathy, gradually regained popularity in the early 2010s due to increasing scientific knowledge and modernization of processing techniques. In 2013, the European Union reauthorized the use of non-ruminant by-products in aquafeeds. Like fishmeal, pig and poultry products are highly digestible, appetizing and well balanced in protein and amino acids. Derived from industrial by-products, they are also less expensive, do not compete with human food sources and have a lower carbon footprint than other ingredients ^(11–13)^. Based on previous studies of salmonids, depending on the percentage of inclusion in aquafeeds, poultry by-products and pig blood meal are suitable replacements for marine ingredients since they do not induce strong adverse effects on fish growth or health ^(14–16)^. However, PAP-based diets effectiveness still requires improvement.

Feed additives, such as yeast and its derivatives, have become of great interest in animal health and disease prevention. In the past few decades, adding yeast to the diets has been well investigated for terrestrial livestock, and positive effects of doing so have been observed for poultry, pigs and ruminants ^(17,18)^. Yeast supplementation has gained popularity for aquaculture species, as positive relations have been identified between nutrition and fish health ^(19)^. As described in the review of Agboola *et al*. ^(20)^, *Saccharomyces cerevisiae* is the yeast strain studied most in aquaculture. It contains relatively high contents of crude protein, amino acids, vitamins and minerals, as well as compounds (MOS and β-glucan) that can stimulate immunity, improve intestinal integrity, assist in wound healing and thus promote growth performances by modulating metabolism ^(21)^. Yeast has been a major ingredient used to mitigate adverse effects of fishmeal-free diets of several fish species and shrimp ^(22– 24)^. Surprisingly, its cytosolic fraction (i.e. yeast extract) is less well known and has been rarely studied, even though it has relatively high contents of protein and bioactive components, such as nucleotides and small peptides, that could also improve fish performances and health ^(25)^.

Due to these qualities and relative lack of use, long-term benefits of a supplementation with the cytosolic fraction of *Saccharomyces cerevisiae* were assessed in this study for rainbow trout fed a fishmeal-free diet composed of poultry by-products and pig blood meal. First, effects on zootechnical parameters were evaluated in a 4-week feed digestibility trial and a 12-week growth experiment. The overall health of the fish was then evaluated, with a focus on intestinal integrity and humoral immunity. Finally, RNAs and miRNAs were sequenced to identify underlying molecular responses of the liver and distal intestine, which are key tissues sensitive to dietary changes.

## 2. Materials and methods

### 2.1. Experimental design

#### Ethical statements

*In vivo* experiments were performed at the French National Research Institute for Agriculture, Food and Environment (INRAE) experimental fish farm facility (IE Numea, 2021 ^(26)^), authorized for animal experimentation by the French Veterinary Service (A64-495-1 and A40-228-1). The experiments were conducted in strict accordance with European Union regulations on the protection of animals used in scientific research (Directive 2010/63/EU) and according to the National Guidelines for Animal Care of the French Ministry of Research (decree no. 2013-118, 02 Jan 2013). In agreement with the ethical committee (C2EA-73), the experiment did not need approval, since it involved only standard rearing practices, with all diets formulated to meet the nutritional requirements (NRC, 2011) of rainbow trout ^(3)^. The staff of the facility received training and personal authorization to perform the experiments.

#### Feeds formulation

To evaluate the benefits of supplementing yeast extract in a fishmeal-free diet, three feeds formulated to be isoproteic, isolipidic and isoenergetic were manufactured by extrusion at INRAE experimental facilities in Donzacq (Landes, France). The main basal composition of the diets differed (Table 1). The control diet (CTL), composed of 19.0% of fishmeal and 7.0% of fish oil, was formulated to approximate a current commercial feed. The processed animal protein (PAP) feed was composed of 10% of dehydrated poultry protein, 6.0% of hydrolyzed feather meal and 1.0% of poultry and pig blood meal. The PAP+YE feed was composed of the PAP feed supplemented with 3% yeast extract with a high content of free amino acids and peptides (Prosaf®, Phileo by Lesaffre, France). The yeast product, derived from the cytosolic fraction of *S. cerevisiae*, was composed of 71.6% crude protein, 8.7% nucleotides, and less than 0.5% lipids (Suppl. Table 1).

**Table 1.** Formulation and proximate composition of experimental diets. CTL, commercial-like diet containing fishmeal and fish oil; PAP, processed animal protein diet; PAP+YE, processed animal protein diet supplemented with 3% yeast extract; DM, dry matter.

|  | CTL | PAP | PAP+YE |
| --- | --- | --- | --- |
| *Ingredient (%)* |  |  |  |
| Corn gluten | 15.0 | 17.9 | 16.0 |
| Soy protein concentrate | 14.0 | 11.0 | 12.0 |
| Soybean meal | 12.0 | 10.5 | 8.3 |
| Faba bean | 8.0 | 12.0 | 8.0 |
| Whole wheat | 10.8 | 9.2 | 13.0 |
| Fishmeal | 19.0 | - | - |
| Dehydrated poultry protein | - | 10.0 | 10.0 |
| Hydrolyzed feather meal | - | 6.0 | 6.0 |
| Poultry and pig blood meal | - | 1.0 | 1.0 |
| Rapeseed oil | 12.2 | 10.5 | 10.5 |
| Fish oil | 7.0 | 7.9 | 7.9 |
| Yeast extract | - | - | 3.0 |
| L-lysine | - | 0.3 | 0.5 |
| L-methionine | - | 0.3 | 0.3 |
| Soy lecithin | - | 0.5 | 0.4 |
| Vitamin premix<sup>1</sup> | 1.0 | 1.0 | 1.0 |
| Dicalcium phosphate | - | 1.0 | 1.0 |
| Mineral premix<sup>2</sup> | 1.0 | 1.0 | 1.0 |
| Yttrium | 0.1 | 0.1 | 0.1 |
| *Proximate composition<sup>3</sup>* |  |  |  |
| Dry matter (%) | 97.1 | 97.4 | 97.2 |
| Protein (%DM) | 46.9 | 46.5 | 46.3 |
| Lipid (%DM) | 22.1 | 21.6 | 20.8 |
| Energy (%DM) | 25.0 | 25.1 | 24.9 |
| Ash (%DM) | 6.76 | 5.53 | 5.59 |
| Starch (%DM) | 12.1 | 13.5 | 14.2 |

<sup>1</sup>Provided per 100 g of premix: vitamin A 500 000 IU, vitamin D3 250 000 IU, vitamin E 5 00 mg, vitamin C 1 429 mg, vitamin B1 10 mg, vitamin B2 50 mg, vitamin B3 100 mg, vitamin B5 200 mg, vitamin B6 30 mg, vitamin B7 3 000 mg, vitamin B8 100 mg, vitamin B9 10 mg, vitamin B12 100 mg, vitamin K3 200 mg, folic acid 10 mg, biotin 100 mg, choline chloride 16 700 mg, and cellulose 76 921mg.
<sup>2</sup>Provided per 100 g of premix: calcium hydrogen phosphate 49 478 mg, calcium carbonate 21 500 mg, sodium chloride 4 000 mg, potassium chloride 9 000 mg, magnesium oxide 12 400 mg, iron sulfate 2 000 mg, zinc sulfate 900 mg, manganese sulfate 300 mg, copper sulfate 300 mg, cobalt chloride 2 mg, potassium iodide 15 mg, sodium selenite 5 mg and sodium fluoride 100 mg.
<sup>3</sup>Analyzed values

#### Digestibility trial

The digestibility trial was conducted at INRAE experimental facilities in Saint Pée-sur-Nivelle (Nouvelle Aquitaine, France). Nine batches of 15 rainbow trout, with a mean body weight (BW) of 100 ± 5 g, were distributed and acclimated into 100 L cylindrical-conical tanks for 15 days. The tanks were supplied with continuous flow (4 L/min) of well-oxygenated fresh water (9 ppm dissolved oxygen) at a constant temperature (17 ± 0.5 °C) under a 12 h photoperiod. Experimental diets, which contained 0.1% yttrium (Thermo Scientific, Waltham, Massachusetts, USA) as an inert marker, were randomly assigned to the tanks (three tanks per diet) (Table 1). Before collecting feces, fish were acclimated to their respective diet for another 15 days period and fed twice a day until visual satiation. During the four weeks following the adaptation period, feces were collected daily using Choubert’s automatic and continuous sieving system ^(27)^, pooled by tank and stored at -20°C until further analytical assays. Apparent digestibility coefficients (ADCs) were determined using the indirect method based on analyzing the yttrium content according to NRC (2011) ^(3)^. Yttrium concentrations were determined in diets and feces after an acid digestion with HNO_3_ and measured with a coupled plasma-mass spectrometry (Agilent 4200 MP-AES, Santa Clara, California, USA). ADCs (%) were estimated for dry matter, energy, protein, lipids, starch, and minerals.

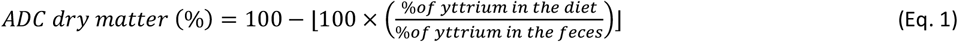

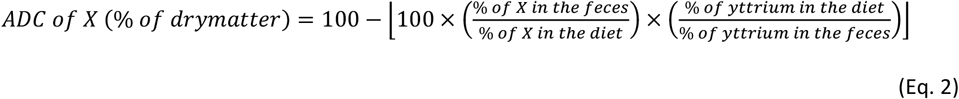

with X the nutrient of interest (*e*.*g*. protein, lipids, starch, minerals) or energy.

#### Growth experiment and sampling procedure

The growth experiment was conducted in the experimental facilities of INRAE in Donzacq (Landes, France). In total, 189 juvenile rainbow trout, with a mean BW of 37 ± 2 g, were divided into groups of 21 fish in 9 tanks of 100 L each supplied with natural freshwater at 17°C. The three experimental diets were randomly allocated to the tanks in triplicate (three tanks per diet) (Table 1). For 12 weeks, fish were reared under a natural photoperiod and fed twice a day by hand to visual satiation. The feed dispensed and total fish biomass in each tank were recorded every three weeks. Zootechnical parameters were calculated for feed efficiency (FE) and daily feed intake (DFI, %/day).

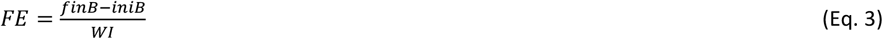

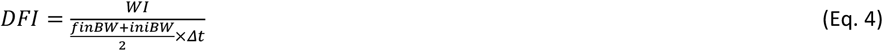

with *finB* and *iniB* as the initial and final biomass per tank (g), respectively; *iniBW* and *finBW* the initial and final BW (g), respectively; *WI* as the feed dispensed during the growth experiment (g); and Δ*t* as the feeding period (days).

After 12 weeks of feeding, fish were anesthetized and euthanized by immersion in successive baths of benzocaine (30 mg/mL and 60 mg/mL, respectively). Nine fish per condition (three per tank) were collected, measured, weighed and frozen at -20°C for analysis of body composition. On these fish, liver and viscera were collected and weighed in order to calculate hepato-somatic index (HSI, %), viscero-somatic index (VSI, %) and condition factor K.

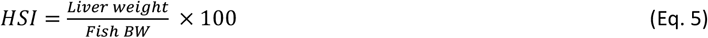

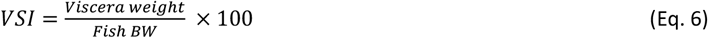

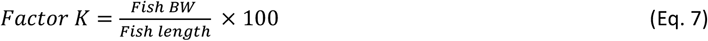

with Fish BW as the fish body weight (g).

An additional 12 fish per condition (four per tank) were anaesthetized and euthanized for tissue sampling. Blood was collected from the caudal vein 48 h after the last meal using a syringe previously treated with EDTA and centrifuged at 3000 x *g* for ten minutes. Plasma was recovered and frozen at -20°C to later quantify humoral immune parameters. Liver and distal intestine samples were collected 24 h after the last meal and soaked in RNA later before being immediately immersed in liquid nitrogen and kept at -80°C until further molecular analysis. Additional samples of proximal and distal intestine were collected 48 h after the last meal, immediately fixed in a 10% buffered formalin solution, and then adequately processed for later histological studies.

### 2.2. Proximate composition of diets, feces and whole body

The proximate compositions of the diets, feces and whole body were quantified according to the following protocols. Dry matter and ashes were determined after 24 hours of drying at 105°C and 7 hours of combustion at 550°C, respectively. Gross energy content was measured using an adiabatic bomb calorimeter (IKA C5003). Crude protein was determined using the Kjeldhal acid-digestion method ^(28)^. The total lipid content was isolated, purified and quantified gravimetrically according to the protocol of Folch *et al*. ^(29)^. Data on the proximate diet and whole-body compositions data were used to calculate protein, lipid and energy intake (kJ/kg BW/day), gain (kJ/kg BW/day) and retention (%), using the following equations:

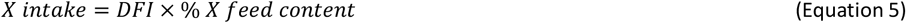

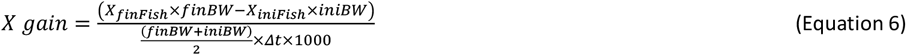

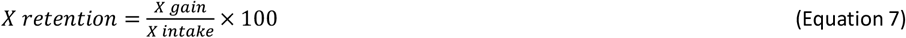

with X as the nutrient considered or energy; X_fin_Fish and X_ini_Fish as the initial and final fish nutrient or energy content, respectively (in % of wet matter).

### 2.3. Humoral immune parameters

Plasma lysozyme activity, peroxidase activity, alternative complement pathway activity and total immunoglobulin M levels were measured in triplicate using a microplate reader (Synergy HT, Biotek, Winooski, Vermont, USA).

#### Plasma lysozyme activity

Plasma lysozyme activity (n = 12 fish per condition) was quantified using a turbidimetric assay, as described by Peixoto *et al*. ^(30)^. On a microplate, 15 µL of plasma were incubated with 250 µL of a *Micrococcus luteus* solution (Sigma-Aldrich, Saint-Louis, Missouri, USA, 0.5 mg/mL in 0.05 M sodium phosphate buffer, pH 6.2) at 25°C. During incubation, absorbance was measured at 0.5 and 15 min at 450 nm. Lyophilized hen egg white lysozyme was serially diluted (0.5 M, pH 6.2) and used to develop a standard curve. Plasma lysozyme concentrations were calculated using the equation for the standard curve.

#### Plasma peroxidase activity

Total plasma peroxidase activity (n = 12 fish per condition) was measured using the protocol of Quade and Roth ^(31)^. Each plasma sample was diluted ten-fold in Ca^2+^- and Mg^2+^-free Hank’s Balanced Salt Solution (HBSS) and incubated 2 min with 50 µL of 3,3′,5,5′-tetramethylbenzidine hydrochloride (TMB, 20 mM) and 50 µL hydrogen peroxide (H_2_O_2_, 5 mM). At the end of incubation, the reaction was stopped by adding 50 µL of sulfuric acid (H_2_SO_4_, 2 M) and the optical density (OD) was read at 450 nm. HBSS was used as a blank and one unit of peroxidase activity was defined as the quantity of peroxidase that produces an absorbance change of 1 OD.

#### Plasma alternative complement pathway activity

Plasma alternative complement pathway activity (n = 9 fish per condition) was estimated using the method of Sunyer and Tort ^(32)^. Rabbit red blood cells (RaRBC) were washed in GVB buffer (isotonic veronal buffered saline with 0.1% gelatin, pH 7.3) and then resuspended in GVB buffer to a concentration of 2.8 × 10^8^ cells/mL. Plasma samples were serially diluted in Mg-EGTA-GVB buffer (GVB buffer containing 10 mM Mg^2+^ and 10 mM EGTA) and incubated in triplicate with 10 µL of RaRBC suspension at room temperature with constant shaking. The hemolysis reaction was stopped after 100 min by adding 150 µL of cold EDTA-GVB (GVB buffer containing 20 mM EDTA). The degree of hemolysis was determined by measuring the OD of the supernatant at 414 nm. Plasma concentrations that yielded 50% of RaRBC hemolysis were defined as ACH50 units. Negative controls consisted of samples without plasma, from which OD values were subtracted for each sample.

#### Plasma total immunoglobulin M level

Total plasma immunoglobulin M levels (n = 12 fish per condition) were determined using an enzyme-linked immunosorbent assay (ELISA) ^(30)^. Plate wells were coated with diluted plasma proteins (1:1000 in 50 mM Na_2_CO_3_ buffer, pH 9.6), blocked 2 h with a blocking buffer (5% low fat milk in TBST, pH 7.3) and washed with a TBST solution (1X tris-buffered saline with 0.1% Tween 20, pH 7.3). Samples were then incubated with 100 µL of anti-rainbow-trout IgM monoclonal antibody (Aquatic Diagnostic Ltd., Scotland, 1:400 in a blocking buffer) for 1 h, washed again with TBST and incubated with an anti-mouse secondary antibody (EMD Millipore Corp., USA, 1:1000 in a blocking buffer) for 1 h. Subsequently, 100 µL of TMB was added, and the reaction was stopped with 100 µL of H_2_SO_4_ (2 M). Absorbance was measured at 450 nm and total immunoglobulin levels were equated to OD.

### 2.4. Histological analysis of the intestine

Proximal and distal intestine samples (n = 12 per condition) were processed using standard histological methods. Briefly, tissues were fixed in 10% buffered formalin before being dehydrated in consecutive ethanol baths (70%, 95% and 100%), clarified in xylene and embedded in paraffin blocks. Subsequently, 2 µm longitudinal sections were cut and stained with hematoxylin and eosin to measure the height of intestinal villi, and additional sections were stained with periodic acid-Schiff alcian blue to quantify the number of goblet cells per mm². Villi height was measured on one slide per fish using a light microscope (Nikon Eclipse E200, Tokyo, Japan) equipped with a digital camera (E3CMOS Series C-mount USB3.0 CMOS Camera, Sony, Tokyo, Japan) and ToupView 3.7 software (ToupTek, Hangzhou, China).

### 2.5. Molecular analysis

#### Total RNA extraction

Total RNA and miRNA was extracted from liver and distal intestine (n = 6 per condition) using a miRNeasy Tissue/Cells Advanced Mini Kit (Qiagen, Hilden, Germany) following the protocol provided by the manufacturer. To avoid genomic DNA contamination, RNA samples (10 µg of each) were subjected to DNase treatment using a Turbo DNA-free kit (Invitrogen, USA). RNA concentrations and quality were determined using a Nanodrop® ND1000 spectrophotometer (Thermo Scientific, USA) and subjected to a Bioanalyzer (Agilent, USA) to validate the RNA extraction procedure. RNA integrity was verified using agarose gel (1%).

#### Preparation and sequencing of cDNA libraries

Total RNA was quantified using a Fragment Analyzer and the standard sensitivity RNA kit (Agilent Technologies, USA). Libraries were constructed using a Stranded mRNA Prep Ligation kit (Illumina, San Diego, California, USA) according to the manufacturer’s instructions. Briefly, poly-A RNAs were purified using oligo(dT) magnetic beads from 500 ng of total RNA. Poly-A RNAs were fragmented and underwent a reverse transcription using random hexamers. During the second strand-generation step, dUTP added dTTP to prevent the second strand from being used as a matrix during the final PCR amplification. Double stranded cDNAs were adenylated at their 3’ ends and ligated to Illumina’s universal anchors. Ligated cDNAs were amplified after 13 PCR cycles. During this PCR, the libraries were indexed, and adapter sequences were added to be compatible with the cluster-generation step. PCR products were purified using AMPure XP Beads (Beckman Coulter Genomics, Brea, California, USA). Libraries were validated using the High Sensitivity NGS kit (Agilent, USA) on the Fragment Analyzer and quantified using the KAPA Library quantification kit (Roche, Basel, Switzerland).

Thirty-six libraries were pooled in equimolar amounts. The pool was then sequenced using a Novaseq 6000 (Illumina, USA) on one lane of an SP flow cell in paired-end 2*150nt mode according to the manufacturer’s instructions.

#### RNA sequencing quality control and transcript quantification

Image analyses and base calling were performed using NovaSeq Control Software and the Real-Time Analysis component (Illumina, USA). Demultiplexing was performed using Illumina’s conversion software (bcl2fastq 2.20). The quality of the raw data was assessed using the FastQC program (version 0.11.8, Babraham Bioinformatics, Cambridge, UK) and the Sequencing Analysis Viewer software (Illumina, USA). The FastQ Screen program (version 0.14, Babraham Bioinformatics, UK) was used to estimate the potential degree of contamination.

Transcripts were quantified using the Nextflow (version 20.11.0-edge) nf-core/rnaseq (revision 3.0) pipeline ^(33)^ on the SLURM cluster of the Genotoul Bioinformatics platform. Nextflow is a FAIR (Findable, Accessible, Interoperable, Reusable) pipeline. The pipeline used the STAR program (version 2.6.1d) ^(34)^ to map the reads to the *O. mykiss* reference genome (Omyk_1.0) and projected the alignments onto the transcriptome (Omyk_1.0.104). The quantification count table was generated using RSEM software (version 1.3.1) ^(35)^.

#### Preparation and sequencing of miRNA libraries

miRNAs were quantified using a Fragment Analyzer and the Small RNA Analysis kit (Agilent, USA). Libraries were constructed using a NEXTFLEX Small RNA-seq v3 kit (Perkin Elmer, Waltham, Massachusetts, USA) following the manufacturer’s instructions. Briefly, a 3’ adenylated adapter was ligated to the 3’ end of 400 ng RNA and purified to remove excess 3’ adapters. A 5’ adapter was ligated to the 5’ end of the 3’-ligated miRNA. The resulting construction was purified to remove excess 5’ adapters. The 5’- and 3’-ligated miRNAs underwent a reverse transcription using an M-MuLV reverse transcriptase and an RT primer complementary to the 3’ adapter. The resulting cDNAs were used as a matrix in a 23-cycle PCR using a pair of uniquely barcoded primers. The resulting barcoded library was size-selected on a Pippin HT using a 3 % agarose cassette (Sage Science, Beverly, Massachusetts, USA), aiming for a size of 147-193 bp. Once size-selected, libraries were verified using a Fragment Analyzer and the High Sensitivity NGS kit (Agilent, USA), and quantified using the KAPA Library quantification kit (Roche, Switzerland).

Thirty-six libraries were pooled in equimolar amounts. The pool was then sequenced using a Novaseq 6000 (Illumina, USA) on one lane of an SP flow cell in paired-end 2*50nt mode according to the manufacturer’s instructions.

#### miRNA sequencing quality control and miRNAs quantification

Quality control of miRNA sequencing was performed as described previously for RNA sequencing. miRNA raw reads were trimmed using the Cutadapt program (version 3.4) ^(36)^ to remove the sequencing adapter (TGGAATTCTCGGGTGCCAAGG) at the 3′ end. Additionally, four bases were also trimmed from the 5′ end and 3′ end of the reads, as described in the manual for the NEXTFLEX Small RNA-Seq v3 kit (Perkin Elmer, USA).

#### miRNA quantification

Trimmed miRNA libraries were processed using the *Prost!* program (version 0.7.60) ^(37)^ to quantify the expression of mature miRNAs. *Prost!* was configured to retain all reads of 17-25 nucleotides that appeared at least five times in the dataset. Sequences were aligned on the USDA_OmykA_1.1 rainbow trout reference genome (accession GCF_013265735.2). Genomic locations were then enriched by *Prost!* using the FishMiRNA database (version of June 2021) and 354 sequences of rainbow trout mature miRNAs described by Cardona *et al*. (2021) ^(38)^. Rainbow trout was used as the main species to annotate the genomic locations, which yielded a “compressed by annotation” table of *Prost!* results that summarized expression values for all known rainbow trout mature miRNAs. As advised by *Prost!* documentation, when multiple potential annotations were identified for a given location, read counts were evenly distributed among the annotations.

#### Analysis of transcripts and miRNAs differential expression

Differential expression was analyzed using the DESeq2 program (version 1.32) ^(39)^, with the RSEM raw counts for total RNA sequencing and the *Prost!* raw expression matrix of annotated mature miRNAs for miRNA sequencing. Differential expression was analyzed in the same way for both datasets using the standard *DEseq2* procedure for all 36 libraries and specifying a suitable contrast for each differential expression test. Genes or mature sequences with a false discovery rate (FDR) less than 0.05 were retained for further analysis.

#### Gene ontology analysis

Enrichment of gene ontology terms was tested in each of the sets of differentially expressed genes. To create a set of background genes, the Manhattan method of the genefilter package ^(40)^ of R software ^(41)^ (version 4.1.3) was used to retrieve a set of 100 genes with expression profiles similar for to those of each differentially expressed genes. Enrichment was then analyzed using the topGO R package (version 2.44.0) ^(42)^ and the Ensembl 104 annotation. All terms of the molecular function, biological process and cellular compartment ontologies were tested using Fisher’s exact test and the “classic” algorithm of topGO.

### 2.6. Statistical analysis of zootechnical parameters, plasma and histological data

Statistical analyses were performed using the Rcmdr R package. Normality of the distributions and homogeneity of the variances were assessed using a Shapiro-Wilk test and Levene’s test, respectively. If normality and homoscedasticity were confirmed, the data were analyzed using one-way or two-way analysis of variance (ANOVA) test followed by Tukey’s range test. If one or both conditions were not met, the data were analyzed using the non-parametric Kruskal-Wallis test. Differences were considered statistically significant at p < 0.05.

## 3. Results

### 3.1. Apparent nutrient digestibility

Overall, ADCs were high for all diets, exceeding 80% for dry matter, protein, lipids, starch and energy, and exceeding 40% for minerals (Table 2). The PAP diet was more digestible than the CTL and PAP+YE diets, with a significant increase in dry matter, protein and energy digestibilities (Tukey’s test, *p* < 0.05). Compared to the digestibilities of the CTL and PAP diets, adding the yeast extract (i.e. PAP+YE) slightly decreased the digestibility of dry matter (by 1.70% and 2.10%, respectively), protein (by 0.44% and 0.55%, respectively) and energy (by 0% and 1.57%, respectively). Nevertheless, all digestibilities remained similar to those of the CTL diet. The digestibilities of lipids, minerals and starch did not differ significantly among the diets.

**Table 2.** Apparent digestibility coefficients (ADCs) of nutrients and energy. CTL, commercial-like diet containing fishmeal and fish oil; PAP, processed animal protein diet; PAP+YE, processed animal protein diet supplemented with 3% yeast extract. Data are presented as the mean ± standard deviation (STD) (n=3 per condition). Degrees of significance (p-value) were estimated with a one-way ANOVA followed by a Tukey’s test, or a Kruskal-Wallis test. Different letters indicate significant differences among groups.

| Nutrients or energy | CTL |  | PAP |  | PAP+YE |  | p-value |
| --- | --- | --- | --- | --- | --- | --- | --- |
|  | Mean | STD | Mean | STD | Mean | STD |  |
| Dry matter | 81.5<sup>c</sup> | 0.24 | 84.6<sup>a</sup> | 0.15 | 82.9<sup>b</sup> | 0.20 | <0.001 |
| Protein | 90.7<sup>b</sup> | 0.42 | 91.6<sup>a</sup> | 0.27 | 91.1<sup>ab</sup> | 0.35 | 0.04 |
| Lipids | 93.8 | 1.89 | 94.5 | 0.48 | 93.6 | 0.05 | 0.56 |
| Minerals | 43.8 | 1.44 | 51.8 | 0.92 | 44.2 | 1.22 | 0.08 |
| Starch | 93.8 | 0.64 | 95.3 | 0.62 | 95.2 | 0.83 | 0.08 |
| Energy | 87.5<sup>b</sup> | 0.63 | 88.9<sup>a</sup> | 0.39 | 87.5<sup>b</sup> | 0.61 | 0.03 |

### 3.2. Growth performances and feed consumption

Fish had the same mean BW at the beginning of the growth experiment (one-way ANOVA, *p* > 0.05), but after 12 weeks of feeding, the group fed the CTL diet showed the best growth performances, with a significantly higher final BW (Table 3). Conversely, as expected, fish fed the PAP diet showed the worst growth performances; however, adding yeast extract to the PAP diet increased fish growth significantly (Tukey’s test, *p* = 0.01). DFI was highest for fish fed the PAP diet (Tukey’s test, *p* < 0.01). The diets did not influence FE, despite a trend for lower FE for fish fed the PAP diet (Kruskal-Wallis, *p* = 0.06). Anatomical indexes, including HSI, VSI and condition factor K, were not affected by the diets (one way ANOVA, *p* > 0.05).

**Table 3.** Growth performances and consumption parameters. CTL, commercial-like diet containing fishmeal and fish oil; PAP, processed animal protein diet; PAP+YE, processed animal protein diet supplemented with 3% yeast extract; SGR, specific growth rate; FE, feed efficiency; DFI, daily feed intake; HSI, hepatosomatic index; VS.I, viscerosomatic index. Data are presented as the mean ± standard deviation (STD) (n=3 per condition). Degrees of significance (p-value) were estimated with a one-way ANOVA followed by a Tukey’s test, or a Kruskal-Wallis test. Different letters indicate significant differences among groups.

|  | CTL |  | PAP |  | PAP+YE |  | p-value |
| --- | --- | --- | --- | --- | --- | --- | --- |
|  | Mean | STD | Mean | STD | Mean | STD |  |
| Initial body weight (g) | 36.7 | 1.98 | 36.6 | 1.00 | 37.1 | 2.32 | 0.99 |
| Final body weight (g) | 230.5<sup>a</sup> | 7.6 | 182.3<sup>c</sup> | 13.5 | 206.7<sup>b</sup> | 14.2 | 0.01 |
| SGR (day<sup>-1</sup>) | 2.19<sup>a</sup> | 0.05 | 1.91<sup>b</sup> | 0.12 | 2.04<sup>ab</sup> | 0.05 | 0.02 |
| FE | 1.16 | 0.01 | 0.99 | 0.04 | 1.15 | 0.03 | 0.06 |
| DFI (%/day) | 1.49<sup>b</sup> | 0.01 | 1.61<sup>a</sup> | 0.04 | 1.44<sup>b</sup> | 0.04 | 0.006 |
| HSI (%) | 1.18 | 0.20 | 1.31 | 0.22 | 1.36 | 0.24 | 0.18 |
| VS.I (%) | 13.7 | 2.10 | 12.97 | 1.39 | 14.09 | 2.19 | 0.47 |
| Condition factor K | 1.65 | 0.14 | 1.62 | 0.13 | 1.58 | 0.17 | 0.67 |

### 3.3. Whole-body composition and nutrients use

At the end of the growth experiment, the diets had not influenced fish body composition of dry matter, ashes, protein, lipids or energy (one-way ANOVA, *p > 0*.*05*) (Table 4). The diets did not affect energy or lipid gains or lipid retention, but protein gain, energy retention and protein retention differed significantly among groups (one-way ANOVA, *p* < 0.05) (Table 5). The control group (CTL) showed the best nutrient utilization capacities for protein gain and protein retention while fish fed the PAP diet had the lowest values for protein gain, energy retention and protein retention. Compared to the PAP diet, the PAP+YE diet increased protein gain (+10.1%), energy retention (+7.0%) and protein retention (+7.3%), with values that did not differ significantly from those of the CTL diet.

**Table 4.** Final whole-body composition. CTL, commercial-like diet containing fishmeal and fish oil; PAP, processed animal protein diet; PAP+YE, processed animal protein diet supplemented with 3% yeast extract; DM, dry matter. Data are presented as the mean ± standard deviation (STD) (n=3 per condition). Degrees of significance (p-value) were estimated with a one-way ANOVA followed by a Tukey’s test, or a Kruskal-Wallis test.

|  | CTL |  | PAP |  | PAP+YE |  | p-value |
| --- | --- | --- | --- | --- | --- | --- | --- |
|  | Mean | STD | Mean | STD | Mean | STD |  |
| Dry matter (%) | 34.1 | 0.2 | 34.5 | 1.3 | 34.6 | 1.7 | 0.73 |
| Ash (% DM) | 5.9 | 0.1 | 5.8 | 0.3 | 5.7 | 0.3 | 0.79 |
| Protein (% DM) | 46.1 | 0.3 | 44.0 | 1.8 | 42.4 | 2.2 | 0.09 |
| Total lipids (% DM) | 51.3 | 0.7 | 55.1 | 2.5 | 57.0 | 7.3 | 0.11 |
| Energy (% DM) | 30.3 | 0.2 | 30.4 | 0.5 | 30.7 | 0.4 | 0.40 |

**Table 5.** Use of the diets. CTL, commercial-like diet containing fishmeal and fish oil; PAP, processed animal protein diet; PAP+YE, processed animal protein diet supplemented with 3% yeast extract. Data are presented as the mean ± standard deviation (STD) (n=3 per condition). Degrees of significance (p-value) were estimated with a one-way ANOVA followed by a Tukey’s test, or a Kruskal-Wallis. Different letters indicate significant differences among groups.

|  | CTL |  | PAP |  | PAP+YE |  | p-value |
| --- | --- | --- | --- | --- | --- | --- | --- |
|  | Mean | STD | Mean | STD | Mean | STD |  |
| Gain |  |  |  |  |  |  |  |
| Energy (kJ/g) | 1.86 | 0.02 | 1.78 | 0.14 | 1.83 | 0.13 | 0.65 |
| Protein (g/kg/day) | 2.73<sup>a</sup> | 0.01 | 2.28<sup>c</sup> | 0.14 | 2.51<sup>b</sup> | 0.04 | 0.002 |
| Lipids (g/kg/day) | 3.18 | 0.02 | 3.39 | 0.70 | 3.38 | 0.36 | 0.67 |
| Retention |  |  |  |  |  |  |  |
| Energy (%) | 51.6<sup>ab</sup> | 0.76 | 45.2<sup>b</sup> | 1.85 | 52.2<sup>a</sup> | 4.07 | 0.03 |
| Protein (%) | 40.4<sup>a</sup> | 0.48 | 31.3<sup>b</sup> | 2.26 | 38.7<sup>a</sup> | 0.93 | 0.03 |
| Lipids (%) | 99.8 | 2.12 | 99.7 | 16.2 | 115.9 | 12.8 | 0.24 |

### 3.4. Histological analysis of the intestine

The villi were significantly higher in the proximal intestine than in the distal intestine (two-way ANOVA, p < 0.01) (Fig. 1). Overall, the diet influenced villi height in both parts of the intestine, with a significant increase in height in fish fed the PAP+YE diet (two-way ANOVA, *p* < 0.05). The number of goblet cells per mm² was significantly larger in the proximal intestine (two-way ANOVA, *p* < 0.001), but the diet had no influence on it (two-way ANOVA, *p* > 0.05) (Fig. 2).

**Fig. 1.**
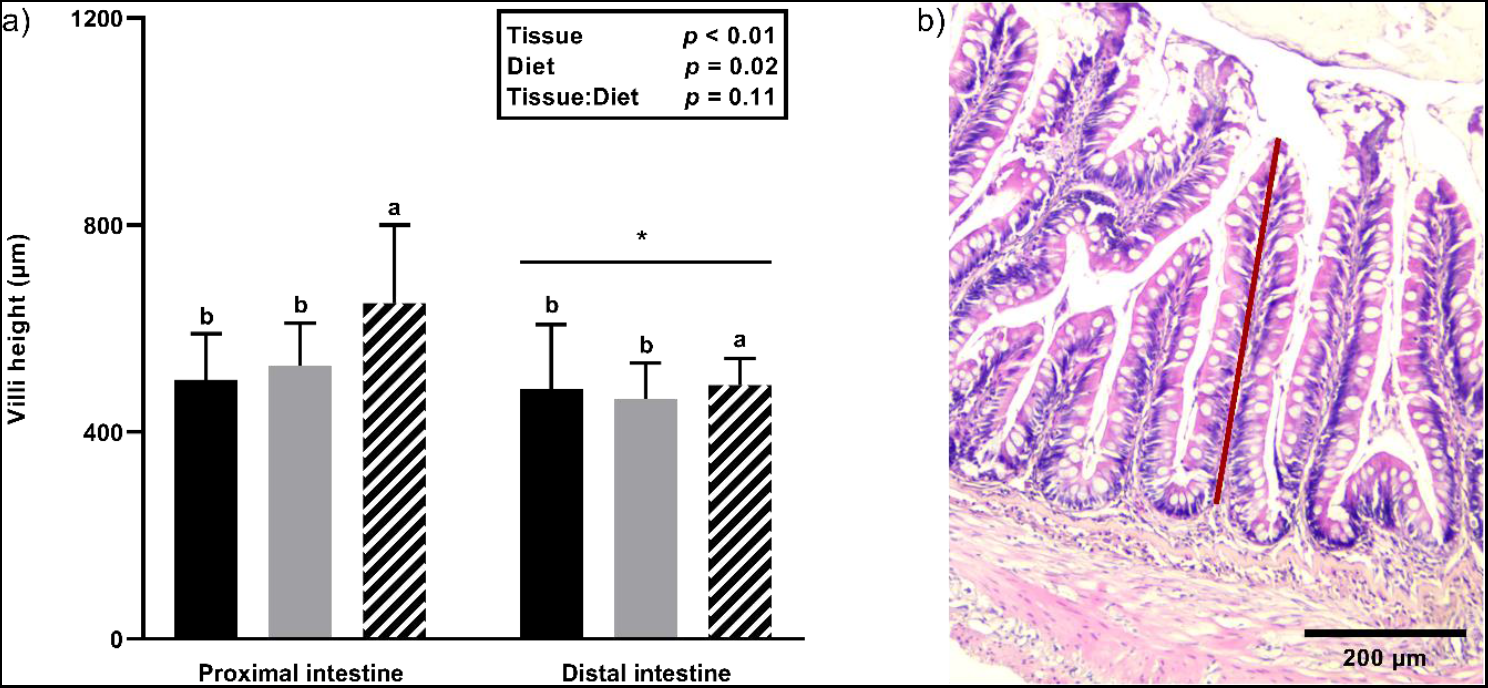
Height of villi in the proximal and distal intestine in rainbow trout after 12 weeks of feeding (a) (■ fish fed the CTL diet; □ fish fed the PAP diet; ▨ fish fed the PAP+YE diet). Villi height was measured from the intestinal epithelium to their apical end, located towards the lumen, as indicated by the red line (b) (n=12 fish per diet). Data are represented as the mean and standard deviation, and were analyzed using a two-way ANOVA. Letters indicate significant differences between diets. * indicates significant differences between tissues. CTL, commercial-like diet containing fishmeal and fish oil; PAP, processed animal protein diet; PAP+YE, processed animal protein feed supplemented with 3% yeast extract.

**Fig. 2.**
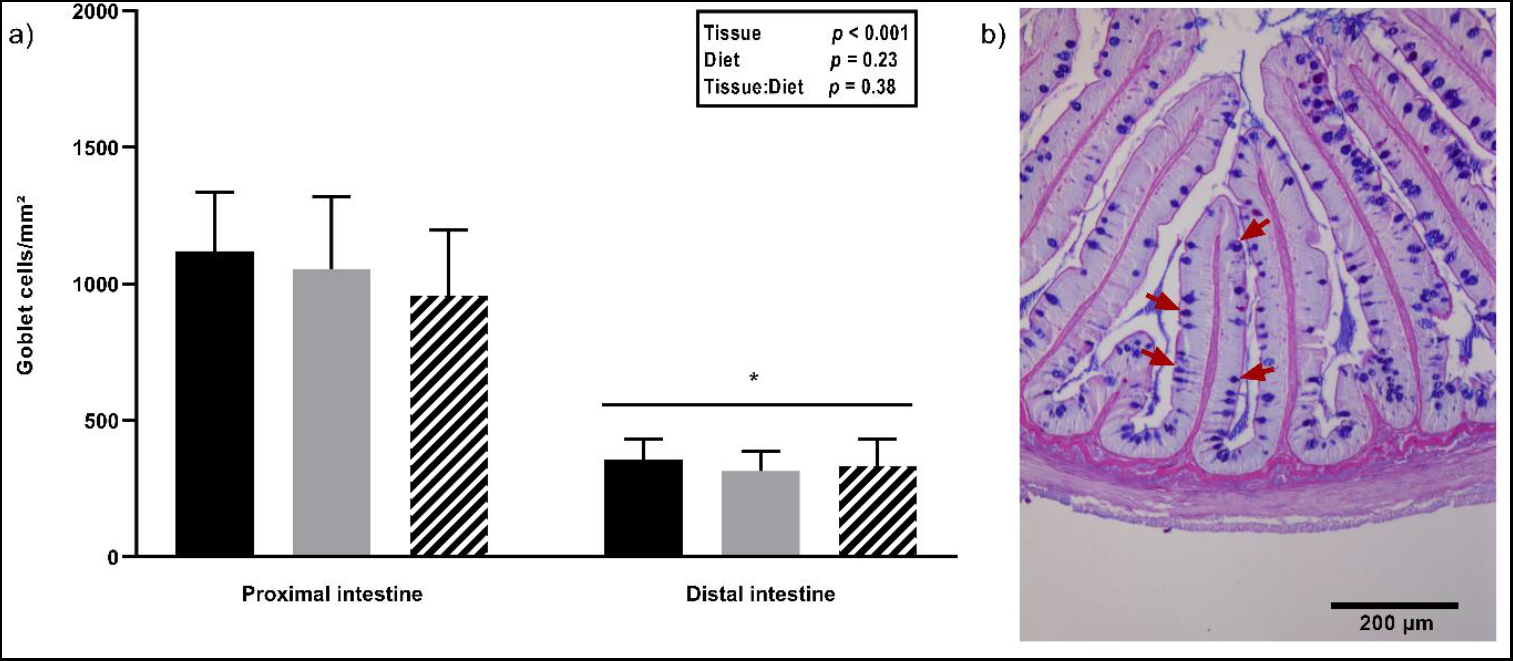
Number of goblet cells per mm² in the proximal and distal intestine in rainbow trout after 12 weeks of feeding (a) (■ fish fed the CTL diet; □ fish fed the PAP diet; ▨ fish fed the PAP+YE diet). Goblet cells are indicated by red arrows and were counted and related to the area of the villi (b) (n=12 fish per diet). Data are represented by as mean and standard deviation, and were analyzed using a two-way ANOVA. * indicates significant differences between tissues. CTL, commercial-like diet containing fishmeal and fish oil; PAP, processed animal protein diet; PAP+YE, processed animal protein feed supplemented with 3% yeast extract.

### 3.5. Plasma immune parameters

The diets did not influence lysozyme and alternative complement pathway activities after 12 weeks of feeding (Table 6). In contrast, total peroxidase activity was significantly lower in fish fed PAP diet than that in fish fed the CTL diet, while that of fish fed the PAP+YE diet was intermediate and also lower that in fish fed the CTL diet (Kruskal-Wallis test, *p* = 0.04). Total immunoglobulin M levels did not differ significantly between the fish fed the CTL and PAP diets but were 25% higher in fish fed the PAP+YE diet (Tukey’s test, *p* = 0.001).

**Table 6.** Plasma immune parameters. CTL, commercial-like diet containing fishmeal and fish oil; PAP, processed animal protein diet; PAP+YE, processed animal protein feed supplemented with 3% yeast extract; ACP, alternative complement pathway. Data are presented as the mean ± standard deviation (STD) (n=9 per condition). Degrees of significance (p-value) were estimated with a one-way ANOVA test or a Kruskal-Wallis. Different letters indicate significant differences among groups.

|  | CTL |  | PAP |  | PAP+YE |  | p-value |
| --- | --- | --- | --- | --- | --- | --- | --- |
|  | Mean | STD | Mean | STD | Mean | STD |  |
| Lysozyme activity ($\mu\text{g/mL}$) | 11.4 | 3.3 | 9.8 | 3.5 | 10.9 | 4.8 | 0.37 |
| Peroxidase activity (U/mL) | 19.7<sup>a</sup> | 17.0 | 9.7<sup>b</sup> | 8.1 | 12.4<sup>ab</sup> | 10.4 | 0.04 |
| ACP (U/mL) | 103.5 | 42.2 | 106.6 | 48.32 | 115.4 | 38.9 | 0.60 |
| Total immunoglobulin (OD<sub>450nm</sub>) | 0.61<sup>b</sup> | 0.18 | 0.70<sup>b</sup> | 0.15 | 0.88<sup>a</sup> | 0.16 | 0.001 |

### 3.6. Analysis of gene expression in the liver and distal intestine

#### RNA sequencing and mapping

SAMtools utilities (version 1.10) ^(43)^ specified that 46.5-93.4 million reads per sample were generated in BAM files, of which 68.0%-83.7% per sample were properly paired. The sequencing produced 18-22 million passed filter clusters per library. The descriptive files as well as the generated data for RNA and miRNAs sequencing are available in the NCBI SRA repository under the following accession number PRJNA924987.

#### Identifying differentially expressed transcripts in the liver and distal intestine in response to the diets, gene ontology enrichment and pathway analysis

In the liver, 14 transcripts were differentially expressed with three down-regulated genes (fold changes ranging from -0.64 to -1.10), and 11 up-regulated genes (fold changes ranging from 0.60 to 2.22) (Fig. 3 a), Table 7). The largest modulation was observed when comparing the PAP and CTL diets, with 11 transcripts differentially expressed, of which one and 10 were down-regulated and up-regulated, respectively. The number of differentially expressed transcripts decreased to three when comparing the PAP and PAP+YE diets, of which two were down-regulated. No genes were differentially expressed when comparing the PAP and CTL diets. Moreover, no transcript was common to all three comparisons. In the distal intestine, 28 transcripts were differentially expressed: 16 transcripts were down-regulated with a range of fold change from -0.44 to -23.67, and 12 were up-regulated with a range of fold change from 0.88 to 22.6 (Fig. 3 b), Table 8). Unlike in the liver, most of them occurred when comparing the PAP+YE and CTL diets, with 13 transcripts, of which six were down-regulated. One of the 13 transcripts was shared with the PAP+YE vs PAP comparison, while two were shared with the PAP vs CTL comparison. The PAP+YE vs PAP comparison identified seven differentially expressed transcripts, of which four where down-regulated. The PAP vs CTL comparison identified eight differentially expressed transcripts, of which two were up-regulated, with one transcript shared with the PAP+YE vs PAP comparison.

**Fig. 3.**
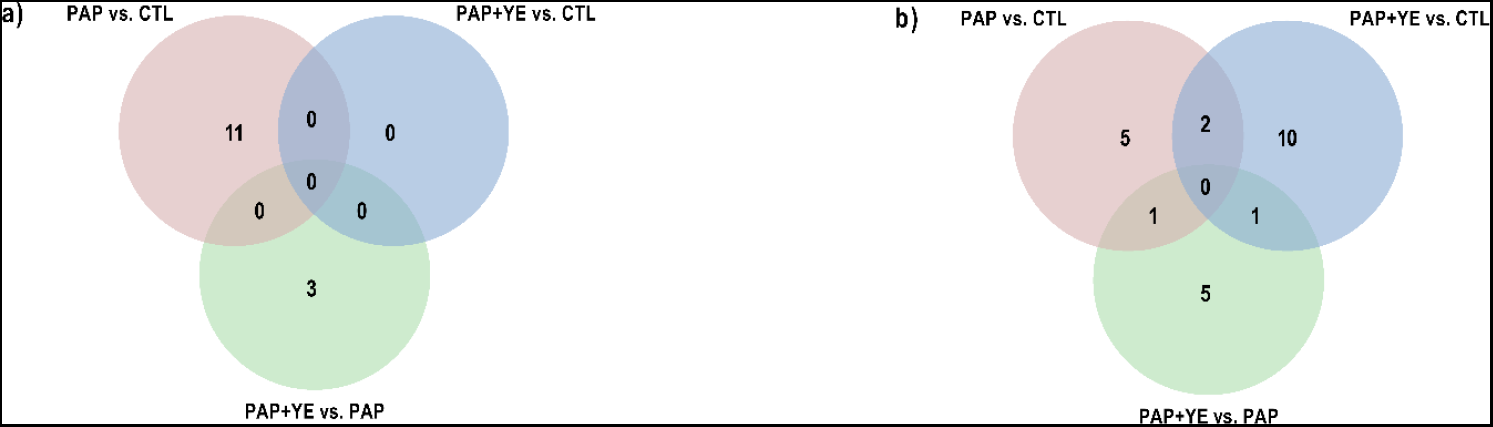
Venn diagrams of RNA sequencing analysis in (a) the liver and (b) distal intestine of rainbow trout fed one of three experimental diets for 12 weeks. Each diagram summarizes the total number of differentially expressed genes in each tissue, the number of differentially expressed genes in each comparison and the common genes (in overlapping regions). CTL, commercial-like diet containing fishmeal and fish oil; PAP, processed animal protein diet; PAP+YE, processed animal protein feed supplemented with 3% yeast extract.

**Table 7.**
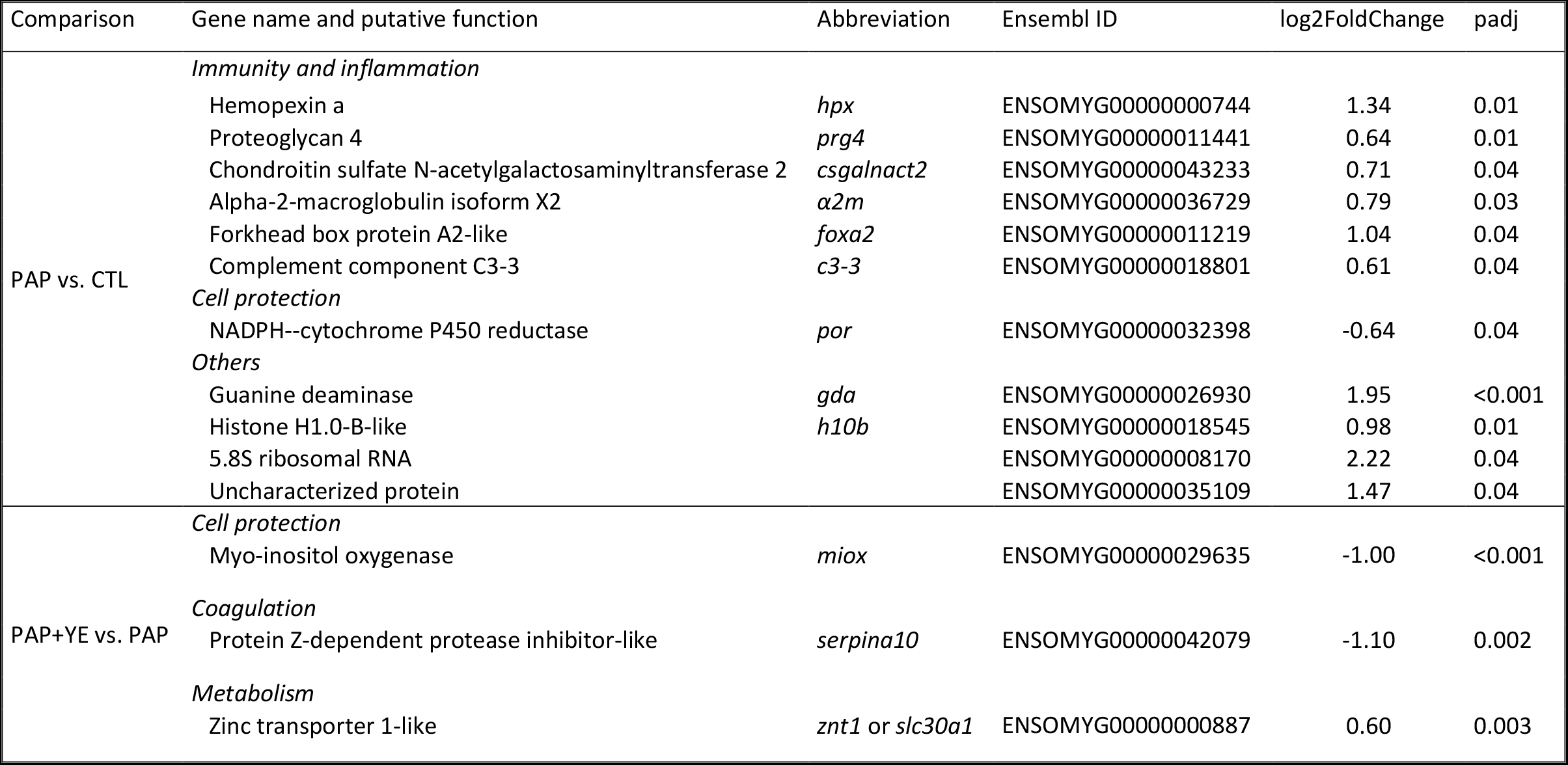
Differentially expressed hepatic genes grouped by diet comparison and putative function. CTL, commercial-like diet containing fishmeal and fish oil; PAP, processed animal protein diet; PAP+YE, processed animal protein diet supplemented with 3% yeast extract. N=6 per condition.

**Table 8.**
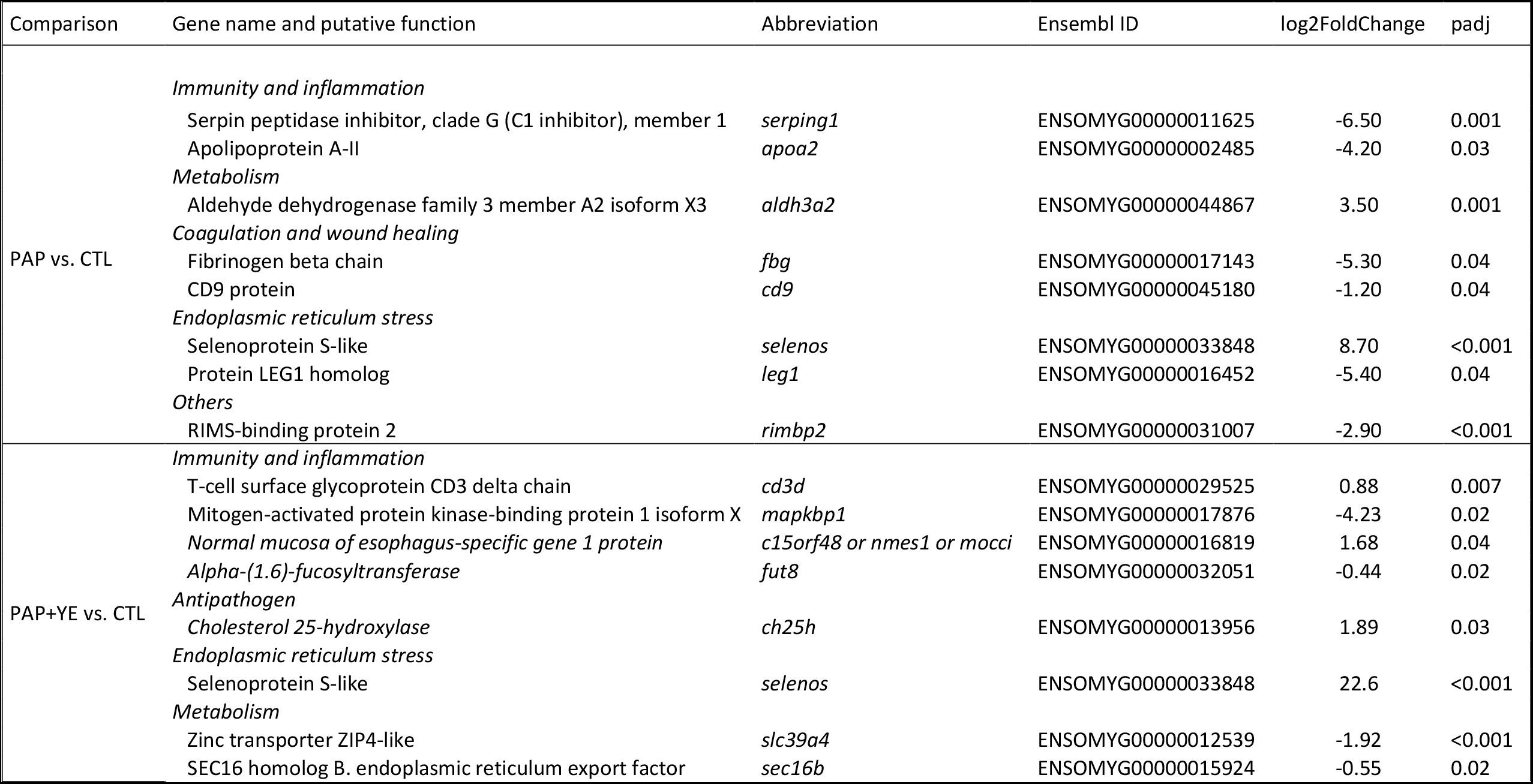
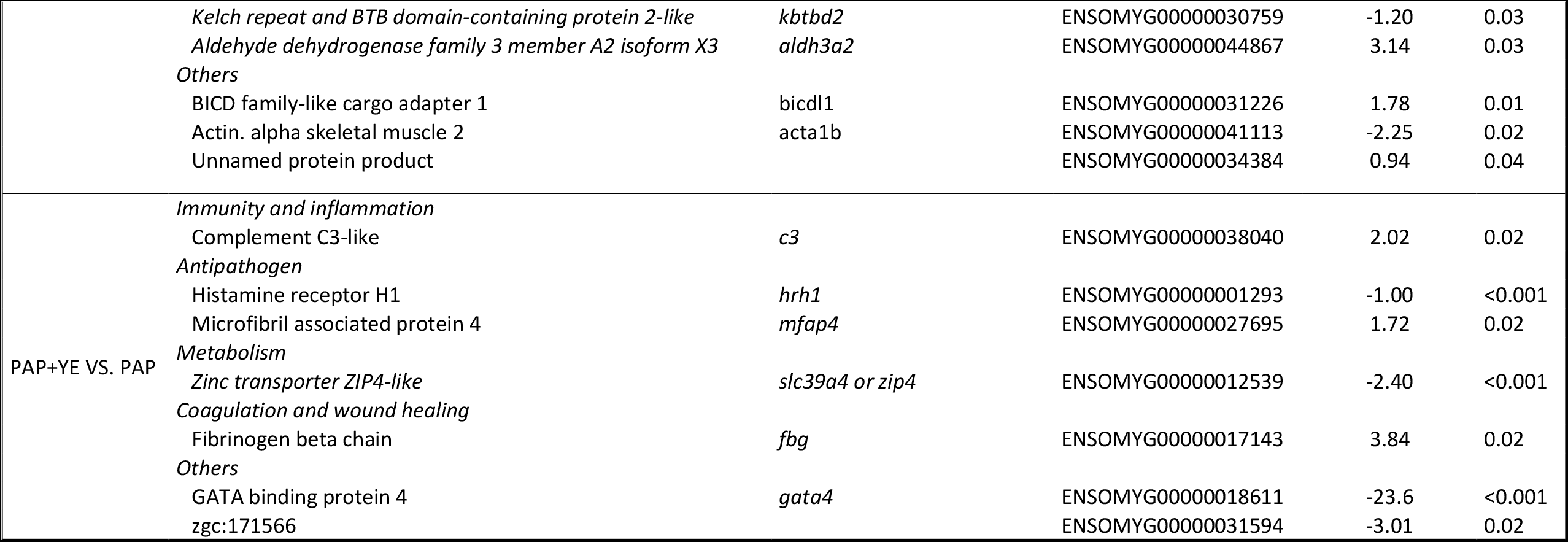
Differentially expressed genes in the distal intestine grouped by diet comparison and putative function. CTL. commercial-like diet containing fishmeal and fish oil; PAP. processed animal protein diet; PAP+YE. processed animal protein diet supplemented with 3% yeast extract. N=6 per condition.

Enrichment of differentially expressed genes was tested against a set of 4990 background genes. Overall, 25-gene ontology terms were found significantly enriched in one contrast. Gene ontology enrichment and an additional literature analysis allowed to potential pathways to be assigned to each differentially expressed transcript (Table 7, Table 8 and Suppl. Fig. 1). In the liver, of the 14 differentially expressed genes, six were related to immune or inflammatory functions. The other eight transcripts were assigned to cell protection, coagulation or metabolism functions. In the distal intestine, of the 28 transcripts differentially expressed, seven were assigned putatively to immunity or inflammation functions, two to coagulation and wound healing, two to endoplasmic reticulum stress response, three to antipathogen functions and two to metabolism. Transcripts classified into the category “*Others*” could not be assigned to specific functions due to lack of literature or poor genome annotation.

### 3.7. Analysis of miRNAs expression in the liver and distal intestine

#### miRNA sequencing and mapping

A total of 321 mature miRNAs with a read count greater than one transcript per million (TPM) were detected in at least one sample. As frequently occurs in miRNA sequencing, the miRNome was dominated by a few highly abundant mature sequences. The most abundant sequence (omy-miR-122-5p) had a mean normalized count of 204 000 TPM across all libraries.

#### miRNAs differentially expressed in the liver and distal intestine in response to the diets

Of the 318 mature miRNAs detected in the liver at more than a mean of 0.1 TPM average, only 4 were differentially expressed, and they belonged to the PAP+YE vs PAP comparison (Table 9). Two were up-regulated (miR-222b-3p and miR-92a-3p) and the other two were down-regulated (miR-18a-5p and miR-217a-5p) in response to the addition of yeast extract. In contrast, the diets had no influence on the differential expression of the 310 mature miRNAs detected in the distal intestine.

**Table 9.**
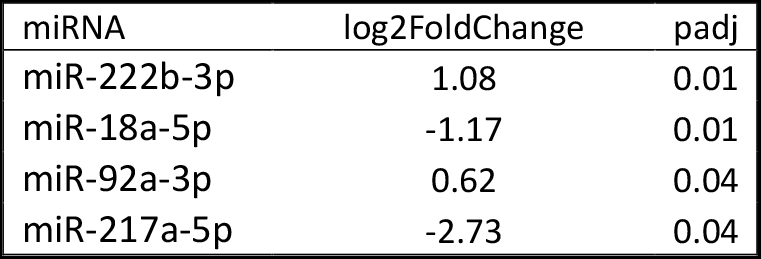
Differentially expressed hepatic miRNAs when comparing the PAP+YE and PAP diets. PAP. processed animal protein diet; PAP+YE. processed animal protein diet supplemented with 3% yeast extract. N=6 per condition.

## 4. Discussion

Replacing fishmeal with alternative ingredients and improving fishmeal-free diets has become increasingly necessary for the sustainable development of aquaculture, especially for carnivorous species of high economic value. It is well established that salmonids, such as rainbow trout, have a low tolerance to fishmeal-free diets. Completely replacing fishmeal with plant-based ingredients is likely to worsen fish performances ^(44,45)^. To counteract the adverse effects on farmed fish growth, alternative animal protein sources (e.g. poultry by-product meal, feather meal, poultry or pig blood meal) have been investigated because of their high protein content. However, results of partially or completely removing fishmeal from the diet remain unsatisfactory for rainbow trout ^(46–48)^, gilthead seabream (*Sparus aurata*) ^(49)^ and Pacific white shrimp (*Litopenaeus vannamei*) ^(50)^. This was confirmed in the present study, which showed that juvenile rainbow trout grow more slowly on a fishmeal-free diet containing plant-based ingredients and PAP than on a 19% fishmeal diet, even though the nutrient requirements were fully met. One solution to improve the performance of fishmeal-free diets is to use functional ingredients such as yeast, which is used not only as a source of protein, but also for its beneficial effects on animal growth and health due to its ability to stimulate the immune system ^(21)^. In the present study, adding 3% yeast extract increased the effectiveness of the PAP diet, as indicated by the final BW of fish fed the PAP+YE diet, which approached that of fish fed the fishmeal-based diet.

Studies agree that animal by-products in diets are appetizing and highly digestible, especially their protein fractions, for several fish species ^(13,51–53)^, which was confirmed in the present study. Because adding yeast to the diet barely changed nutrient digestibility, the increased growth of fish fed the PAP+YE diet was due not to an increase in digestibility, but to better metabolic use of nutrients. Indeed, adding yeast tended to increase the FE, and trout fed the PAP+YE diet retained more protein than those fed the PAP diet. Because rainbow trout are more tolerant of soybean meal than Atlantic salmon are ^(54)^., the small decrease in soybean meal content of the PAP+YE diet was likely not the main cause of the increase in growth. The benefits of yeast for growth have been demonstrated for many fish species such as salmon ^(55)^ and tilapia (*Oreochromis mossambicus*) ^(56)^. However, these studies used yeast to replace fishmeal protein and thus included it in percentages higher than the 3% used in the present study. The protein in the yeast extract (71.6 %DM) used in the present was composed mainly of small peptides and free amino acids. This characteristic may have increased the bioavailability of amino acids, as previously demonstrated for rainbow trout ^(57)^, and thus promoted amino acid use and growth. The increased growth could also have been related to a positive effect of the yeast extract on the structure of the intestinal mucosa. Trout fed the PAP+YE diet had significantly higher intestinal villi, which agrees with previous observations of Eurasian carp (*Cyprinus carpio*) fed a diet in which fishmeal was partially replaced with 3% of a complex product derived from the enzymatic hydrolysis of yeast ^(58)^. These results suggest that adding yeast extract does not harm the intestinal mucosa but instead provides a larger surface area for nutrient absorption, particularly for amino acids and peptides, due to its positive effect on the height of villi ^(59)^.

Strong evidence suggests a relation among fish nutrition, health and growth performances. In this respect, plasma immune markers and gut structure were analyzed extensively, since they have been identified as relevant for assessing fish health ^(60,61)^. An unbiased transcriptomic analysis was performed on genes and miRNAs in the liver and distal intestine. In the plasma, only total peroxidase activity was significantly lower in fish fed the PAP diet, and the diets did not influence lysozyme or alternative complement pathway activities. In the gut, enteritis is usually indicated by an increase in the number of goblet cells and shortening of the intestinal villi ^(62,63)^. This phenotype, which is found in Atlantic salmon fed a diet rich in soybean products ^(64)^, was not observed in the present study, in which the mean number of goblet cells was identical among the three diets. Fish fed the PAP and PAP+YE diets did not have shorter villi than those fed the CTL diet. Therefore, plasma immune indicators and intestinal histology did not indicate an exacerbated inflammatory-type immune response in fish fed diets containing animal by-products, with or without yeast. Nevertheless, fish fed the supplemented PAP+YE diet had the highest level of total immunoglobulin M, which can be explained by the high content of small peptides (50.1% of total peptides are smaller than 0.50 kDa) in the yeast extract ^(65)^. Peptides from a variety of sources are known to have nutraceutical effects due to their antioxidant, antimicrobial and immunomodulatory properties ^(66)^. Regarding their influence on immunity, bioactive peptides can stimulate lymphocytes and thus the secretion of immunoglobulins ^(67,68)^, which has been reported for large yellow croakers (*Pseudosciaena crocea R*.) fed a diet supplemented with a fish-protein hydrolysate assumed to contain bioactive peptides ^(59)^. Thus, our results suggest that supplementation with a peptide-rich yeast extract could improve acquired humoral immunity and consequently fish defenses against pathogenic stresses. This hypothesis will need to be verified through a pathogenic challenge.

Unlike previous studies that performed high-throughput sequencing transcriptomic analysis of fish, including rainbow trout ^(69,70)^, few genes were differentially expressed in the liver and distal intestine. This difference could be related to the timing of sampling, which occurred 24 h after the last meal. Thus, the modulations were likely related more to background changes in fish physiology than to short-term postprandial effects. While the PAP diet altered the transcriptomic profile of the liver, adding only 3% yeast extract to it appeared to restore a hepatic physiological response similar to that obtained with the CTL diet, which is consistent with the observed phenotypes. The transcriptomic response of the gut to the diets was less distinct, although most of the differentially expressed genes were observed in fish fed the PAP+YE or CTL diet. While analysis of plasma immune markers and intestinal histology studies did not provide clear evidence of inflammation, as confirmed by the lack of induction of key cytokines in the gut immune response, analysis of gene ontology terms and the literature revealed that a large proportion of the differentially expressed transcripts were related to inflammation and immune functions. In the liver, nearly half of the differentially expressed transcripts were assigned putatively to immune or inflammatory functions, and induced in fish fed the PAP diet compared to those fed the CTL diet. Specifically, the analysis revealed the up-regulation of two acute phase proteins (*hpx* ^(71)^ and *α2m* ^(72)^), as well as a transcriptional regulator of acute-phase responses (*foxa2* ^(73,74)^). The protein encoded by *prg4*, which is responsible for activating and stimulating hepatic macrophages in rats ^(75)^, and the enzyme encoded by *csgalnact2*, which is required for elongation during chondroitin sulfate synthesis, were also up-regulated in fish fed the PAP diet. In the distal intestine, effects associated with immune functions were prevalent in trout fed the PAP+YE diet, suggesting that the cytosolic fraction of yeast could have specifically targeted immune- or inflammation-related genes of the intestinal mucosa (i.e. *serping1* ^(76)^, *apoa2* ^(77)^, *mapk1b* ^(78)^, and *c15orf48* ^(79)^). Also, two of them were related to stimulation of T- and/or B-cells (i.e. *fut8* ^(80,81)^ and *cd3d* ^(82)^). In the liver and distal intestine, the C3 component was the only complement member that was differentially expressed among the diet. The complement system plays a fundamental role in the innate immune system since it is involved in phagocytosis, inflammatory reactions and production of antibodies. Among the many proteins involved in activating the complement cascade, the C3 molecule is one of the most important ^(83)^. In the present study, the dietary regulations differed between these two tissues. While the liver was more sensitive to the PAP diet, the distal intestine was more sensitive to the PAP+YE diet. These results are consistent with an increase in expression of intestinal complement C3 in response to the use of yeast in gilthead seabream ^(84,85)^. In fish fed the diet PAP+YE, the transcriptomic response of the intestine also revealed the modulation of genes related to mechanisms of pathogen control or allergic reaction. Indeed, we observed an up-regulation of *ch25h*, the key enzyme in the production of 25-hydroxycholesterol, a powerful soluble antiviral factor that restricts the propagation of viruses ^(86)^ and that was induced in response to viral infections in Atlantic salmon ^(87)^. Compared to the PAP diet, the PAP+YE diet modulated expression of antipathogen-related genes *mfap4* and *hrh1*. In Nile tilapia (*Oreochromis niloticus*), MFAP4 proteins are known to agglutinate and opsonize bacterial pathogens, which helps to prevent bacterial infections ^(88)^. The histamine receptor 1, encoded by the *hrh1* gene, is an effector of the action of histamine on phagocyte function in mammals and fish ^(89,90)^. Overall, our results indicate that including yeast in the diet of rainbow trout could strengthen defense capacities of the gut, which is one of the main contact and entry points of pathogens and allergens in the body. Of all the differentially expressed genes, selenoprotein S-like (*selenos*) varies the most in response to diet, and its expression is highly pronounced in the intestine in response to diets that contain terrestrial animal by-products. Selenos is a protein in the endoplasmic reticulum that is involved in reducing oxidative and endoplasmic reticulum stress ^(91)^. The issue of whether PAP diets induce oxidative and endoplasmic reticulum stress remains to be investigated, since the transcriptomic analysis revealed no marker genes for stress that would support this hypothesis. We also observed the modulation of certain genes putatively involved in coagulation and healing in both the liver and distal intestine. In the intestine, fish fed the PAP diet showed a down-regulation of coagulation markers *fbg* ^(92)^ and *cd9* ^(93)^, suggesting a reduction in coagulation capacity. However, adding 3% yeast extract may have restored it, as indicated by the opposite effect of the PAP+YE diet on the expression of *fbg* in the intestine and a downregulation of anticoagulant *serpina10* ^(94)^ in the liver.

The analysis of small non-coding RNAs did not help to identify mechanisms that underlay the observed growth phenotypes. Under the study conditions (samples collected 24 hours after the last meal), few miRNAs were modulated in response to the diet (only four in the liver and none in the distal intestine). However, miR-217-5p varied the most in expression, which in mammals appears to target inflammatory mechanisms ^(95)^. Its expression decreased greatly in response to the PAP+YE diet.

In conclusion, the study confirms the utility of adding yeast products to improve the nutritional quality of fishmeal-free diets for rainbow trout. The mechanisms that underlay the decreased growth performance of fish fed a diet without fishmeal remain unclear. The results showed no consistent inflammatory reaction, but they did suggest that the molecular response of fish to the fishmeal-free diets stimulates both innate and acquired defense mechanisms, including those that fight pathogens and allergens, as well as those that support tissue coagulation and healing processes. Supplementation with yeast extract could induce some of these mechanisms, which confirms that yeast products could increase the robustness and growth of rainbow trout. In animal nutrition, yeast is considered beneficial for growth and health. The yeast extract used to supplement the PAP diet was the cytosolic fraction, rich in small peptides, free amino acids and some nucleotides. Thus, the observed effects on growth, intestinal histology and gene expression were likely due to the nucleotides, or the presence of specific bioactive peptides, which should be investigated in future studies.

## 5. Acknowledgment

We dedicate this study to Frank Sandres, who cared for the fish during this study and sadly passed away. We thank Anthony Lanuque for monitoring the growth experiment. We are grateful to the Genotoul bioinformatics platform Toulouse Occitanie (Bioinfo Genotoul, https://doi.org/10.15454/1.5572369328961167E12) for providing help, computing resources and storage resources. We would also like to thank Christophe Klopp for his advices in the analysis of the high-throughput sequencing data.

## 6. Financial support

This study was financially supported by the iSIte E2S-UPPA project. Diogo Peixoto and Benjamin Costas were supported by FCT -Fundação para a Ciência e a Tecnologia, Portugal (UI/BD/150900/2021 and 2020.00290.CEECIND, respectively). This study also received funds from FCT within the scope of UIDB/04423/2020 and UIDP/04423/2020 granted to CIIMAR.

## 7. Conflict of Interest

None.

## 8. Authorship

Study design: SSC, NR, KP

Funding acquisition: SSC

Animal experiments: FT, PA

Feed formulation: FT, NR, SSC, KP, LF

Biological analyses: LF, DP, CT

Bioinformatics analyses: LF, SMH, CG, CK, JB

Statistical analysis: LF, CG, BC

Project administration and supervision: SSC, NR, KP

Writing - original draft: LF

Writing - review & editing: LF, SSC, KP, NR, BC, JB, DP, CT, CG, SMH, CK

---

**Supplementary Table 1.** Analytical composition (% dry matter (DM)) of the yeast extract (Prosaf^®^, Phileo by Lesaffre).

| Component | Analyzed valyes |
| --- | --- |
| Dry matter (%) | 95.7 |
| Crude protein (% DM) | 71.6 |
| Crude lipid (% DM) | < 0.5 |
| Ash (% DM) | 8.9 |
| Energy (kJ/g DM) | 20.1 |
| Total nucleotides (% DM) | 8.7 |
| Phosphorus (g/kg DM) | 13.8 |
| Arginine (% DM) | 3.1 |
| Histidine (% DM) | 1.3 |
| Isoleucine (% DM) | 2.8 |
| Leucine (% DM) | 4.0 |
| Lysine (% DM) | 4.6 |
| Threonine (% DM) | 2.7 |
| Valine (% DM) | 3.6 |
| Methionine (% DM) | 0.9 |
| Cysteine (% DM) | 0.8 |
| Phenylalanine (% DM) | 2.3 |
| Tyrosine (% DM) | 2.0 |
| Tryptophan (% DM) | 0.8 |
| Alanine (% DM) | 4.5 |
| Aspartic acid (% DM) | 6.4 |
| Glutamic acid (% DM) | 12.0 |
| Glycine (% DM) | 2.8 |
| Proline (% DM) | 2.9 |
| Serine (% DM) | 2.7 |

## Notes

### Competing Interest Statement

The authors have declared no competing interest.

